# The Leray-XT COI primer pair is not suitable for observing ciliates and radiolarians

**DOI:** 10.64898/2026.04.30.721870

**Authors:** Isabelle Ewers, Quentin Mauvisseau, Mahwash Jamy, Sonja Rückert, Frédéric Mahé, Micah Dunthorn

## Abstract

The Leray-XT primer pair has been widely used to amplify the mitochondrial cytochrome c oxidase subunit I (COI) gene from animals. In some marine metabarcoding studies, protists have also been amplified and sequenced using these primers. Here, we ask if the Leray-XT COI primer pair is suitable for observing ciliates and radiolarians, which are numerically and ecologically important components of marine protistan communities. We show that while there are sufficient COI reference sequences for ciliates in NCBI for taxonomic assignments, there are currently only two COI reference sequences for radiolarians. Using *in-silico* analyses, we additionally show that while the reverse primer Leray-XT primer can bind and potentially amplify both ciliates and radiolarians, the forward primer cannot bind to either taxon. These results show that the Leray-XT primer pair is not suitable for observing ciliates and radiolarians, although it may be useful for observing other marine protistan taxa.

## Introduction

Metabarcoding is a powerful approach to observe protistan communities in marine environments (de Vargas et al. 2015; Piredda et al. 2018; Wang et al. 2020; Markussen Bjorbækmo et al. 2023; Ramond et al. 2023; Santoferrara et al. 2023). This method is only powerful, though, if there are available reference sequences that can be used to taxonomically assign metabarcoding data, and if the chosen primers can amplify taxonomic groups of interest (Taberlet et al. 2018; Santoferrara et al. 2020). Parts of the small subunit of the rRNA (SSU-rRNA) gene are commonly targeted in environmental metabarcoding studies of protists because there are curated reference databases (Guillou et al. 2013; del Campo et al. 2018; Berney et al. 2022) and because a lot of SSU-rRNA primers are able to amplify many of the major protistan taxa (Pawlowski et al. 2012; Vaulot et al. 2022). Marine protistan communities were recently revealed in metabarcoding studies that used the Leray-XT primer pair (Wangensteen et al. 2018; Bakker et al. 2019; Turon et al. 2022). However, it is not yet known if the Leray-XT primers, which amplify a fragment of the mitochondrial cytochrome c oxidase subunit I gene (COI), are useful for observing abundant marine protistan taxa such as ciliates and radiolarians.

Ciliates and radiolarians are important components of microbial eukaryotic communities in most marine ecosystems (de Vargas et al. 2015; Keeling and del Campo 2017; Giner et al. 2020; Santoferrara et al. 2020; Burki et al. 2021; Laget et al. 2024). Ciliates are morphologically diverse protists that are important bacterial grazers (Lynn 2008; Dolan et al. 2013; Weisse and Montagnes 2022), although some can be mixotrophic (Esteban et al. 2010). Radiolarians are equally diverse protists that are involved in numerous marine biogeochemical processes such as the silica and carbon cycles (Decelle et al. 2015; Suzuki and Not 2015; Biard 2022).

Leray et al. (2013) designed what became known as the Leray primer pair to amplify COI from marine metazoans. The Leray primer pair’s performance was originally tested using genomic DNA from 30 animal phyla, as well as on environmental DNA extracted from fish guts collected in French Polynesia (Leray et al. 2013). Wangensteen et al. (2018) subsequently modified and increased the degeneracy of the Leray primer pair to develop what became known as the Leray-XT primer pair. The performance of the Leray-XT primer pair was originally tested on environmental DNA extracted from samples collected from hard-bottom marine communities in the Cíes and Balearic Islands (Wangensteen et al. 2018). While metazoans were targeted in that metabarcoding study, Wangensteen et al. (2018) also uncovered very few ciliates and no radiolarians in their data, although they did find numerous other protistan groups such as red algae, diatoms, and dinoflagellates. Using the same Leray-XT primer pair, Bakker et al. (2019) metabarcoded pelagic samples from the Caribbean Sea, and Turon et al. (2022) metabarcoded samples from inside and outside salmonid aquaculture cages in Norway. These two studies also found few ciliates and no radiolarians.

Here we evaluated if the Leray-XT primer pair is suitable to observe ciliates and radiolarians in environmental metabarcoding studies. We first evaluated the number of COI sequences deposited in NCBI’s Nucleotide Database and Sequence Read Archive (SRA) that were identified as being derived from either a ciliate or a radiolarian. We then used *in-silico* analyses to evaluate the ability of the Leray-XT primer pair to amplify a part of the COI gene region of these two protistan groups. Results from this study can be used to interpret the diversity of protistan communities as estimated from environmental metabarcoding using the Leray-XT primer pair.

## Methods

### Searching NCBI for available COI sequences

On 18 March 2026, ciliates and radiolarians were searched in NCBI’s (Sayers et al. 2020) taxonomy browser to determine their taxon ID. The NCBI taxon ID was then used to determine how many sequences are available specifically for the COI gene by searching for the respective taxon ID (e.g., “txid2763[Organism:exp]” for Ciliophora) in combination with the query “AND (COI OR COX1 OR CO1 OR cytochrome c oxidase subunit 1) NOT (COX2 OR COX3)” or slight variations of that query, when required. The filter “Protists” in the nucleotide database was activated during all searches. Since Radiolaria is not listed as a taxon in NCBI’s taxonomy and consequently does not have a taxon ID, the three radiolarian subgroups Acantharea, Polycystinea, and Sticholonchida were individually used with their respective taxon IDs. NCBI’s sequence read archive and genome database were also searched for mitochondrial genomes assigned to ciliates or radiolarians using the taxon IDs in combination with the query “AND (mitochondrial genome OR mitochondrion)”. We did not evaluate COI reference databases (Ratnasingham and Hebert 2007; González-Miguéns et al. 2024), as these are subsets of what is available on NCBI.

### Searching for binding sites in available COI sequences

The Leray-XT primer pair consists of the 26 base pair long forward primer mlCOIintF-XT (5′-GGWACWRGWTGRACWITITAYCCYCC-3′), modified by Wangensteen et al. (2018) from the forward primer mlCOIintF (5’-GGWACWGGWTGAACWGTWTAYCCYCC-3’) designed by Leray et al. (2013), and of the 26 base pair long reverse primer jgHCO2198 (5’-TAIACYTCIGGRTGICCRAARAAYCA-3’) designed by Geller et al. (2013). To find out whether the Leray-XT primer pair can potentially bind, and therefore amplify, the COI gene region of the identified COI or mitochondrial genome sequences of the taxa from above, a subset of those sequences was selected and downloaded in FASTA format. The primer binding sites in each sequence were examined to determine whether they match with the sequence of the Leray-XT primer pair using Bioedit v.7.2.5 (Hall 1999).

Despite many COI sequences being available for the ciliates on NCBI, the primer evaluation was conducted using ten mitochondrial genome sequences of ten different ciliate species. This was due to a lot of the COI sequence entries being replicates, or them not including the binding sites for the Leray-XT primer pair, making them unsuitable for the in-silico analyses. From each of the selected mitochondrial genome sequences, the COI gene region was extracted and, together with the eleven selected COI sequences, a ClustalW (Thompson et al. 1994) multiple sequence alignment was generated in Bioedit. To evaluate the Leray-XT forward primer, the primer sequence was searched in each sequence taking into account the degenerate nucleotide positions. To search for the binding site of the Leray-XT reverse primer, the reverse complement of the reverse primer sequence was used for the evaluation, also considering the degenerate nucleotide positions. To check the findings of the above analyses, a visual alignment of the Leray-XT primer pair and DNA sequences of the previously mentioned protistan taxa was generated using the Geneious Prime v2025.1.2 (Kearse et al. 2012). The DNA sequences were imported from NCBI using the “NCBI Nucleotide” option and the primers were manually added using the “New Sequence” option. The DNA sequences and Leray-XT primer pair were aligned using a “Geneious Alignment” with the “Global alignment with free end gaps” and “51% similarity (5.0/-3.0)” options. A visual assessment of mismatches on the binding sites was then performed.

## Results and Discussion

### Ciliates

There are currently 3,384 COI sequences deposited in NCBI’s Nucleotide Database, and there are 40 COI sequences and mitochondrial genome entries in the Sequence Read Archive (**Table S1**). Most of these ciliate sequences are derived from barcoding efforts of terrestrial and freshwater species (Chantangsi et al. 2007; Strüder-Kypke and Lynn 2010; Greczek-Stachura et al. 2021), although some are from marine species (Jung et al. 2018; Moon et al. 2019; Park et al. 2019; Hu et al. 2022). A few sequences were from genomes of terrestrial ciliates deposited in GenBank (Pritchard et al. 1990; Burger et al. 2000; de Graaf et al. 2009; Berendonk 2011), which are not easily searchable when just looking for COI. Part of the reason that so few sequences are available for ciliates is that there is not yet a COI primer pair that can amplify all ciliate subtaxa (Strüder-Kypke and Lynn 2010), and ciliate barcoding efforts have shifted to the D1-D2 region of the large subunit of the ribosomal rDNA locus (Pawlowski et al. 2012; Santoferrara et al. 2013; Stoeck et al. 2014; Abraham et al. 2019). Although there are not as many ciliate COI sequences deposited in NCBI as compared to protistan groups such as the red algae (data not shown), there are likely enough available references from enough ciliate subtaxa that can be used to taxonomically assign COI metabarcoding sequences as being derived from a ciliate, but maybe not be able to assign them to a specific genus or higher subtaxon.

The *in-silico* analyses showed that while the reverse primer of the Leray-XT primer pair has matching binding sites in ciliates, the forward primer has no such matching binding sites (**Figs S1**,**S2**). The COI gene sequence of the 9 different ciliate species showed 7 to 9 mismatches to occur between the forward primer and the supposed binding site in the DNA sequence. Most of these mismatches occur in the 3’ and 5’ end regions (i.e., the first and last 5 nucleotides) of the forward primer, which is likely to affect the stability of the primer binding and can lead to PCR failure. These effects can be exacerbated when mismatches are present in the 3’ end region (Stadhouders et al. 2010). That is, the Leray-XT primer pair cannot bind to ciliates and therefore cannot amplify this large and important group of marine protists. This finding calls into question the observation of ciliates when using this primer pair by Wangensteen et al. (2018), Bakker et al. (2019), and Turon et al. (2022), as these studies should not have resulted in successful amplifications. These studies used ciliate COI sequences from BOLD (Ratnasingham and Hebert 2007) as taxonomic references, but some of the ciliate reference sequences were misidentified and now are labelled as “Heterokontophyta” (Wangensteen pers. comm.). This finding also supports the findings of Strüder-Kypke and Lynn (2010) that we still lack a universal primer pair to amplify COI in ciliates.

### Radiolarians

There are currently no COI sequences deposited in NCBI’s Nucleotide Database, although there are two COI sequences in single cell genome entries in the Sequence Read Archive (**Table S1**). These are the genomes from the marine *Acanthometra* sp. and *Lithomelissa* sp. (Macher et al. 2023). Barcoding efforts of the radiolarians have instead focused on sequencing the nuclear small subunit rRNA (Pawlowski et al. 2012; Biard et al. 2017; Sandin et al. 2025). Because there are only two radiolarian COI sequences deposited in NCBI (which are part of single-cell genome sequences deposited in the Sequence Read Archive), there are likely not enough available references that can be used to taxonomically assign all COI metabarcoding sequences derived from radiolarians, with many sequences likely being unassignable.

The *in-silico* analyses showed that while the reverse primer of the Leray-XT primer pair has matching binding sites in radiolarians, the forward primer has limited matching binding sites, as 4 mismatches occur between forward primer and the DNA sequence of *Acanthometra* sp. and 5 mismatches occur between forward primer and the DNA sequence of *Lithomelissa* sp (**Figs S1**,**S2**). Like with the ciliates, these mismatches occur in the 3’ and 5’ end regions of the forward primer and are therefore likely to affect the stability of the primer binding. Therefore, when amplifying DNA of radiolarians using the Leray-XT forward primer, the mismatches will likely prevent the primer from properly binding to the target DNA, leading to the Radiolaria not being amplified and recovered at all. Yet, even in the case of successful amplification (e.g., in the case of lower annealing temperatures during PCR), the lack of available COI reference sequences in online databases prevents the detection and identification of radiolarians in environmental samples for now. This finding supports the lack of observation of radiolarians when using this primer pair by Wangensteen et al. (2018), Bakker et al. (2019), and Turon et al. (2022), as these studies did not result in successful amplifications and identifications.

It should be noted that, in contrast to the ciliates, for which many reference sequences and mitochondrial genomes are available that could be included in the primer evaluation analyses, the same analyses could only be performed on two taxa of the Radiolaria. Thus, it remains to be uncovered whether other radiolarian taxa have the same or different numbers of mismatches with the Leray-XT forward primer or whether these mismatches are present at the same nucleotide positions. It could therefore be possible that the forward primer is able to bind to and amplify the DNA of other taxa in this even larger and arguably more important group of marine protists.

### Efficacy of the Leray-XT primer pair for observing ciliates and radiolarians

Widely-used SSU-rRNA primers are known to be unable to observe some protistan groups (Vaulot et al. 2022), but they are still employed to draw general conclusions about protistan biodiversity and biogeography (de Vargas et al. 2015; Piredda et al. 2018; Wang et al. 2020; Markussen Bjorbækmo et al. 2023; Ramond et al. 2023; Santoferrara et al. 2023). The Leray-XT primer pair can likewise be used to draw general conclusions about marine protistan biodiversity and biogeography. It is important to note, though, that the numerically and ecologically important ciliates and radiolarians will not be observed because of their inability to be amplified with this primer pair that originally designed to observe marine metazoans and, in the case of the radiolarians, there are not enough available sequences in the reference databases.

## Acknowledgments

We thank Bánk Beszteri, Florian Leese, and Owen Wangensteen for helpful comments. Thanks also go to Sam Carter and Taylor Swift for providing background music.

## Data Availability Statement

All data that support the findings of this study are available in the main text or Supplementary Information.

---

**Table S1:**
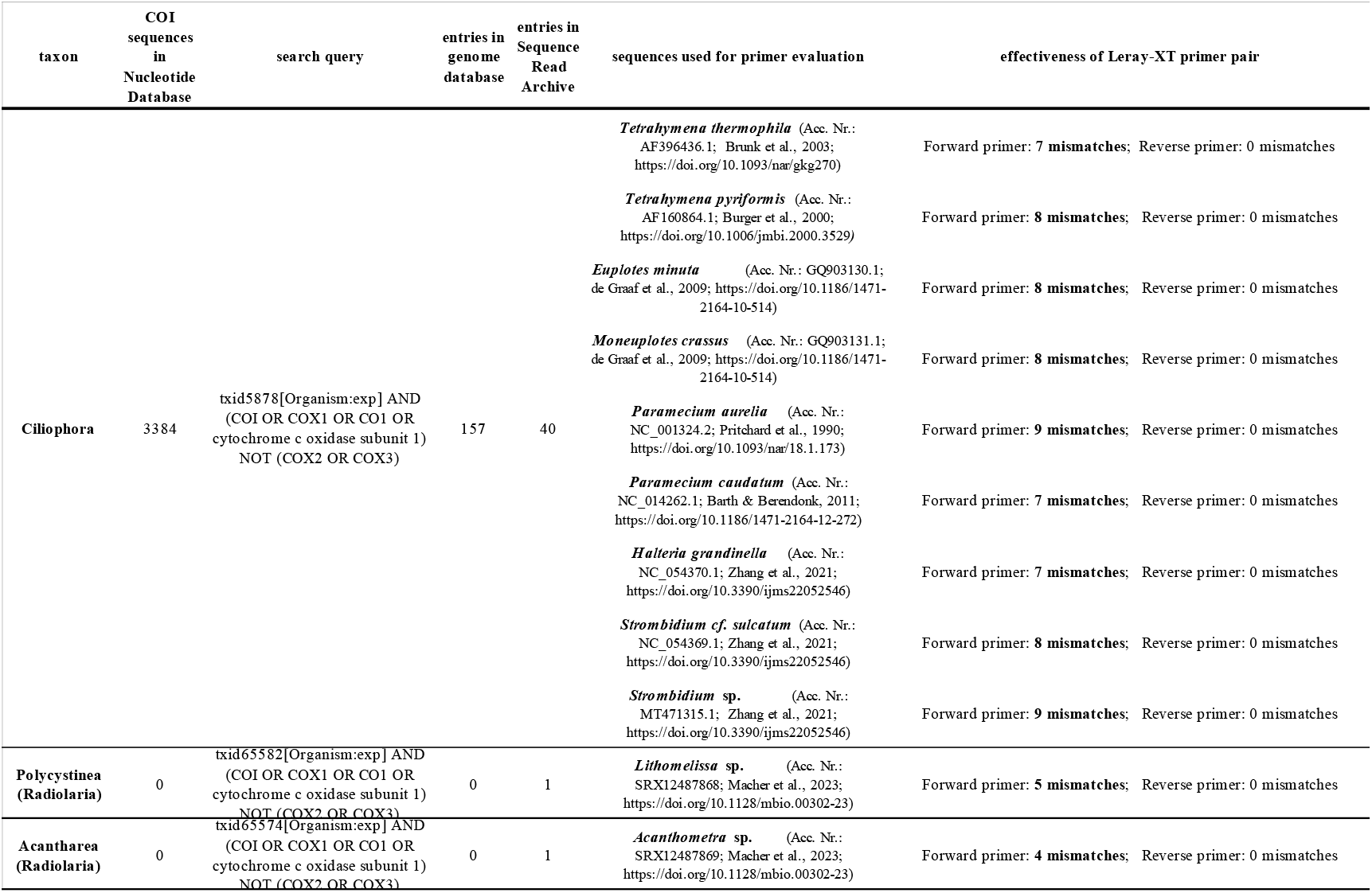
Table showing the available number of COI reference sequences in NCBI’s nucleotide database, as well as the number of entries in NCBI’s genome database and in the Sequence Read Archive for the ciliates and the three groups of the Radiolaria. Further listed are the taxa and their DNA sequences (accession numbers) used for the Leray-XT primer pair evaluation, as well as the effectiveness of the primer to amplify these protist groups in terms of mismatches.

**Figure S1:**
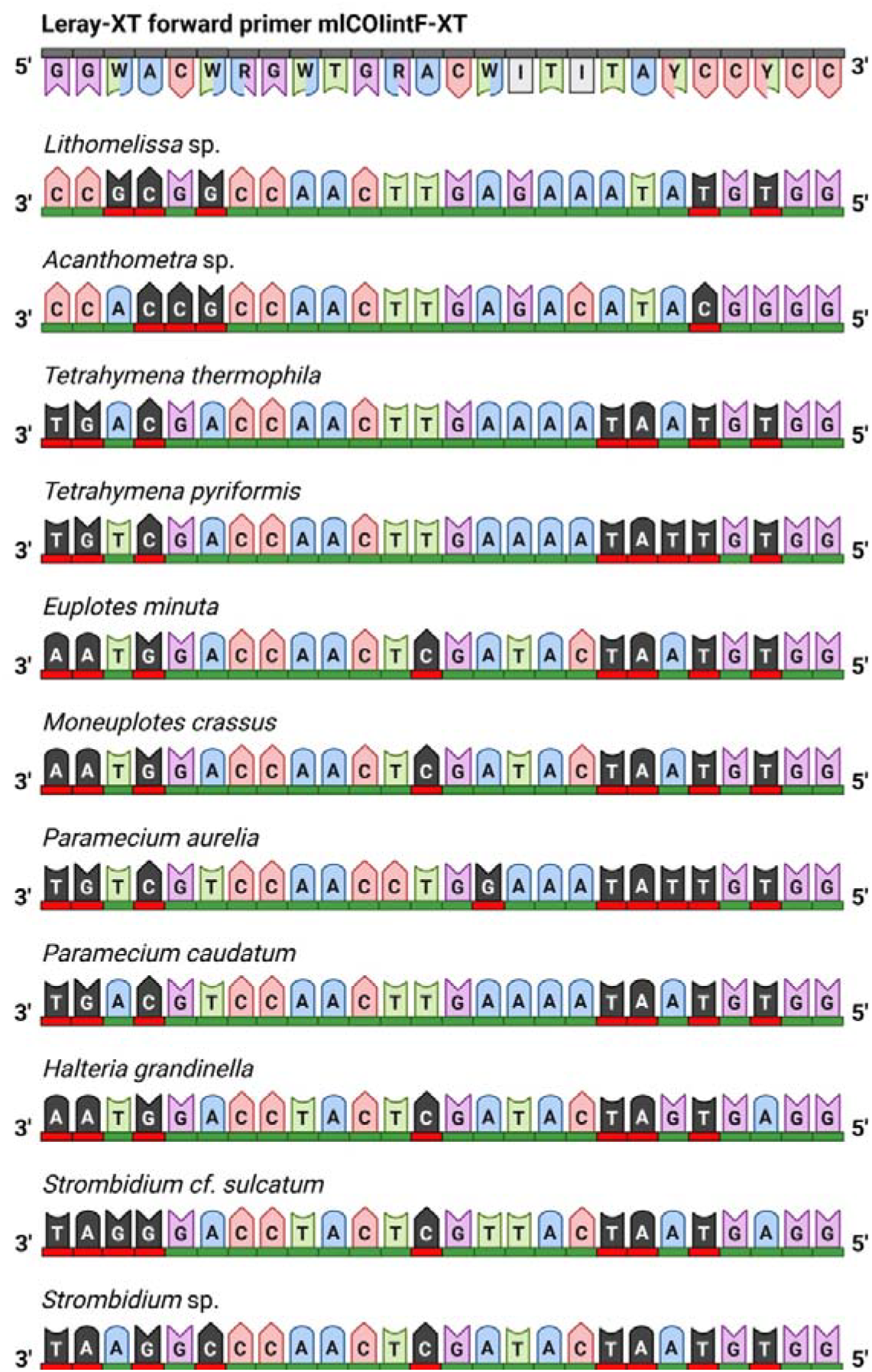
Ability of the Leray-XT forward primer to bind to the DNA sequences of two species of radiolarians and nine species of ciliates for evaluation of primer effectiveness to detect both protist groups in environmental DNA samples. Red and black nucleotides indicate mismatches between primer sequence and target DNA sequence.

**Figure S2:**
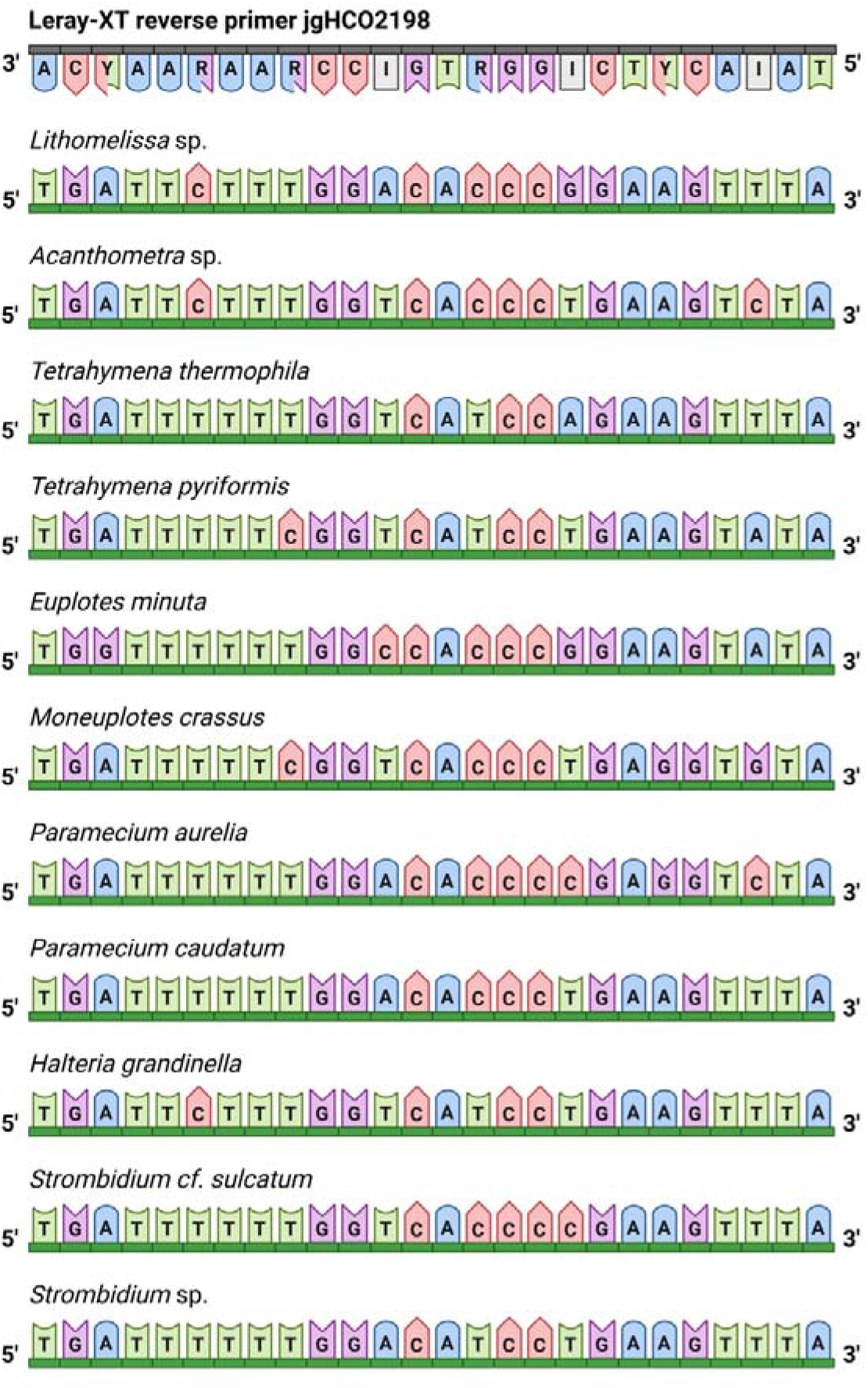
Ability of the Leray-XT reverse primer to bind to the DNA sequences of two species of radiolarians and 9 species of ciliates for evaluation of primer effectiveness to detect both protist groups.

